# Imaging-based screen identifies novel natural compounds that perturb cell and chloroplast division in *Chlamydomonas reinhardtii*

**DOI:** 10.1101/2024.12.30.630779

**Authors:** Manuella R. Clark-Cotton, Sheng-An Chen, Aracely Gomez, Aditya Mulabagal, Adriana Perry, Varenyam Malhotra, Masayuki Onishi

**Author notes:** Address correspondence to: Masayuki Onishi. These authors contributed as co-first authors to this work (Manuella R. Clark-Cotton, Sheng-An Chen).

## Abstract

Successful cell division requires faithful division and segregation of organelles into daughter cells. The unicellular alga *Chlamydomonas reinhardtii* has a single, large chloroplast whose division is spatiotemporally coordinated with furrowing. Cytoskeletal structures form in the same plane at the midzone of the dividing chloroplast (FtsZ) and the cell (microtubules), but how these structures are coordinated is not understood. Previous work showed that loss of F-actin blocks chloroplast division but not furrow ingression, suggesting that pharmacological perturbations can disorganize these events. In this study, we developed an imaging platform to screen natural compounds that perturb cell division while monitoring FtsZ and microtubules and identified 70 unique compounds. One compound, curcumin, has been proposed to bind to both FtsZ and tubulin proteins in bacteria and eukaryotes, respectively. In *C. reinhardtii,* where both targets coexist and are involved in cell division, curcumin at a specific dose range caused a severe disruption of the FtsZ ring in chloroplast while leaving the furrow-associated microtubule structures largely intact. Time-lapse imaging showed that loss of FtsZ and chloroplast division failure delayed the completion of furrowing but not the initiation, suggesting that the chloroplast-division checkpoint proposed in other algae requires FtsZ or is absent altogether in *C. reinhardtii*.

**SIGNIFICANCE STATEMENT:**

- Successful cell division requires the coordination of both organelle inheritance and cytokinesis. The unicellular alga *Chlamydomonas reinhardtii*, which spatiotemporally coordinates the division of its chloroplast with cytokinesis, is an excellent model to study the regulation.
- We screened libraries of natural compounds for perturbations of cell and/or chloroplast division, identifying 70 unique chemicals. By time-lapse microscopy using one of the hits, curcumin, we demonstrate that although chloroplast division failures delay the completion of cytokinesis, it does not impair initiation.
- These findings suggest that the chloroplast-division checkpoint proposed in other algae requires FtsZ or is absent altogether in *C. reinhardtii*.

## INTRODUCTION

In the majority of eukaryotes, cytokinesis is achieved by ingression of the plasma membrane to form a cleavage furrow. The best-studied model of cleavage-furrow formation is the contractile actomyosin ring model (Schroeder, 1973; Fujiwara and Pollard, 1976; Pollard and O’Shaughnessy, 2019), which posits that the contractile force generated by filamentous actin (F-actin) and myosin II drives membrane ingression. Despite the widespread adoption of the model, taxa other than Opisthokonts (animals, fungi, and related species) and Amoebozoa lack myosin II (Odronitz and Kollmar, 2007), limiting the contractile actomyosin ring model’s explanatory power to a small fraction of eukaryotes. Therefore, a fuller understanding of eukaryotic cytokinesis requires an evolutionary approach that includes a diversity of organisms.

Successful cell proliferation and differentiation require fine coordination of cytokinesis with division and segregation of other components of the cell. The coordination of mitosis and chromosome segregation with cytokinesis is the subject of decades of research (Cullati and Gould, 2019; Holder *et al*., 2019; Vavrdová *et al*., 2019). Multiple studies have also uncovered the spatiotemporal regulation of organelle distribution in dividing cells, such as for the ER, Golgi, peroxisomes, and mitochondria (Jongsma *et al*., 2015; Mascanzoni *et al*., 2019). One of the lesser-studied organelles for their coordination with cell division is the chloroplast found in plants and eukaryotic algae. Chloroplasts in the vast majority of photosynthetic eukaryotes derive from the primary endosymbiosis event, in which an early eukaryotic ancestor engulfed and established a stable endosymbiosis with a cyanobacterium (McFadden, 1999). The chloroplast division apparatus consists of the inner FtsZ ring of bacterial origin and the outer dynamin-like ring of eukaryotic origin (Mori *et al*., 2001; Vitha *et al*., 2001; Kuroiwa *et al*., 2002). While the cell-cycle regulated expression of these proteins is well studied (Sumiya *et al*., 2016), how they are positioned relative to the cell-division plane and how their function is regulated during cytokinesis remain poorly understood. Importantly, many unicellular algae have a single chloroplast per cell, and some land plants have cell types with few chloroplasts (de Vries and Gould, 2018; MacLeod *et al*., 2022), making such coordination vital for their proliferation and function.

Natural products are valuable tools for investigating cytokinesis. Large-scale screens for division-perturbing compounds have been performed in various species, including yeast (Dunstan *et al*., 2002) and animal cells (Shoemaker, 2006). However, the range of organisms typically used for such screens overlaps with the distribution of myosin II, limiting the evolutionary scope of the insights this work can reveal. Additionally, such large-scale screens have not been conducted using photosynthetic organisms.

The unicellular alga *Chlamydomonas reinhardtii* is an emerging model of cell division without myosin II (Cross and Umen, 2015; Breker *et al*., 2016; Breker *et al*., 2018; Onishi *et al*., 2020). Furthermore, in *C. reinhardtii*, a single chloroplast lies in the cell division plane, and its partition is spatiotemporally coordinated with cell division (Goodenough, 1970). Recent work revealed that loss of F-actin by treatment with the natural compound and cytoskeletal toxin Latrunculin B (LatB) can dramatically impair chloroplast division, suggesting F-actin’s role as a positive regulator of this coordination (Onishi *et al*., 2020).

In this study, we describe the results of a microscopy-based chemical-genetic screen and report several previously unreported natural compounds that perturb cell proliferation in *C. reinhardtii*. Using time-lapse imaging, we also demonstrate how cytoskeletal structures are altered during division failures.

## RESULTS AND DISCUSSION

### Live-cell markers for monitoring mitosis, cytokinesis, and chloroplast division

To precisely monitor the effects of natural compounds on cell division and chloroplast division, we developed a live-cell imaging platform utilizing fluorescently tagged markers. EB1 is a microtubule plus-end binding protein (Pedersen *et al*., 2003) that decorates the growing tips of microtubules. In *C. reinhardtii*, EB1-mNeonGreen (mNG) has been used to monitor dynamic cortical microtubules in interphase cells, the mitotic spindle, and the furrow-associated microtubules (Harris *et al*., 2016; Onishi *et al*., 2020).

The *C. reinhardtii* genome encodes two FtsZ homologs, *FTSZ1* and *FTSZ2*, both amenable to tagging with fluorescent proteins (this study) and allows for the monitoring of chloroplast division.

We performed time-lapse imaging of cells expressing EB1-mScarlet (mSc) and FtsZ2-mNG. As expected, EB1-mSc labeled the mitotic spindle and the cleavage furrow during cell division (Figure 1A; Movie S1). Shortly before the mitotic spindle appeared, FtsZ2-mNG localized to the chloroplast midzone, forming a ring-like structure perpendicular to the mitotic spindle. After mitosis, EB1-mSc decorated the ingressing cleavage furrow (Onishi *et al*., 2020) (Figure 1A); Movie S1, while the FtsZ2-mNG ring contracted concomitantly with chloroplast division (Figure 1A; Movie S1).

**FIGURE 1:**
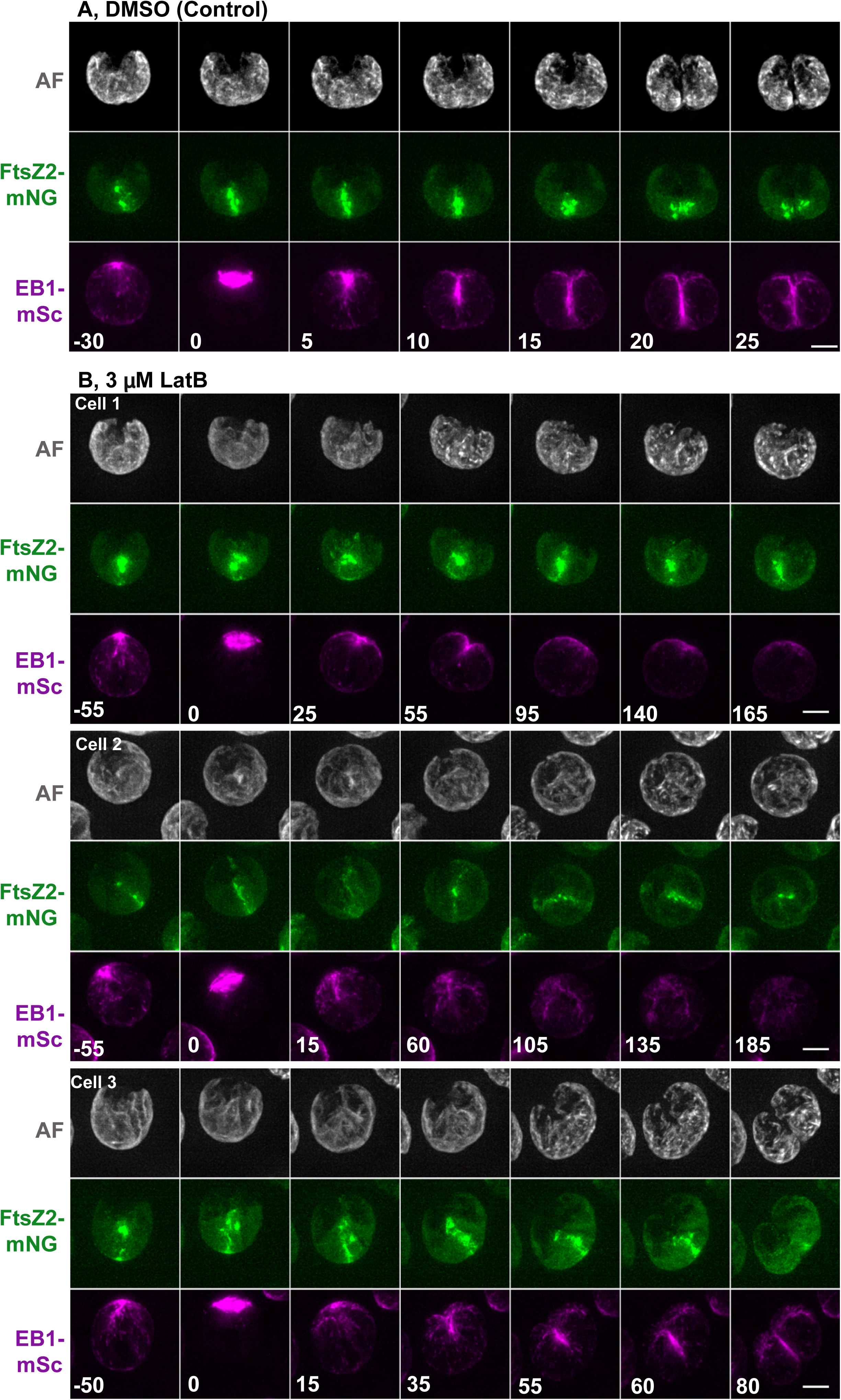
LatB treatment perturbs the FtsZ2 ring and impairs the coordination of cell and chloroplast division. (A) In DMSO (0.03%)-treated *nap1-1* cells, an FtsZ2-mNG ring formed at the chloroplast midzone in preprophase, and EB1-mSc marked the advancing furrow in the dividing cell. The cell and chloroplast divided within 20 min of spindle appearance. (B) Three types of division abnormalities were observed in *nap1-1* cells treated with 3 µM LatB: (Cell 1) A fragmented FtsZ2-mNG midzone structure appeared in preprophase, and the furrow is transiently formed (frame 4) but did not advance. (Cell 2) A weak FtsZ2-mNG ring-like structure was formed, and no furrow was observed; (Cell 3) A fragmented FtsZ2-mNG signal converged into a midzone structure (frame 3), and a furrow ingressed slowly across the cell, seemingly cleaving through the undivided chloroplast. For (A and B), time is set to zero when a spindle is observed and the first timeframe is selected based on the initial appearance of FtsZ2-mNG signal. For (B), select time-points are shown to capture events during the slow division process in these cells. Strain, SCC022 (*FtsZ2-mNG EB1-mSc nap1-1*). Scale bars = 5 μm.

### Effects of latrunculin B on cell and chloroplast division

To confirm that our imaging platform is capable of detecting defects caused by drug treatment, we used a test drug, LatB. Previous report showed that disruption of F-actin structures dramatically impairs chloroplast division, while the cleavage furrow can still form as long as the drug is added after the cell has reached a critical size (Onishi *et al*., 2020). We reasoned that LatB might altering the formation, placement, and/or stability of the FtsZ ring. To test this, we generated an EB1-mSc FtsZ2-mNG tagged strain in the *nap1-1* background, which lacks the divergent and latrunculin-resistant actin NAP1 and therefore is sensitive to the drug treatment (Onishi *et al*., 2016; Onishi *et al*., 2018).

Indeed, we observed three types of ring abnormality. First, some FtsZ2 structures appeared as a fragmented ring (Figure 1B, Cell 1; Movie S2), which remained concentrated at the midzone as a poorly defined structure. The beginning of a furrow was briefly observed (frames 3-4), but the furrow eventually retracted. Second, FtsZ2-mNG structures were ring-like but not limited to the midzone, appearing at different regions of the chloroplast over time while the EB1-mSc signal remained stable (Figure 1B, Cell 2; Movie S3). Finally, we observed cells in which a FtsZ2-mNG structure appeared at the midzone, accompanied by a similarly oriented cleavage furrow (Figure 1B, Cell 3; Movie S4). Although the FtsZ2-mNG structure appears patchier than in untreated cells, this orientation of FtsZ2-mNG ring and cleavage furrow supported both cell and chloroplast division, albeit requiring approximately three times as long as in untreated cells. Our imaging platform allowed for the detection of defects in cytoskeletal structures and cell and chloroplast division caused by LatB. Thus, it might enable the identification of novel compounds that perturb cytoskeletal structures, chloroplast division, and/or cell division.

### Previously unreported natural compounds impair cell and chloroplast division in *C. reinhardtii*

To broadly probe for compounds that affect cell and/or chloroplast division, we screened the National Cancer Institute’s Natural Products Set V (NCI; 390 compounds) and the TimTec Natural Product Library (TT; 720 compounds), which together include diverse bacterial, fungal, plant, and animal sources. We used the *nap1-1* strain background to allow for the possible identification of actin-perturbing compounds whose effects might be obscured by NAP1. *EB1-mSc FtsZ2-mNG nap1-1* cells synchronized to late-G1/S phase were added to 96-well plates pre-populated with compounds (Figure 2; final concentration was 25 µM), and the cells were visually inspected after 3 hours, when untreated control cells had completed division, identifying 72 hits. Several fields were imaged for each hit to capture defects in cell division (differential interference contrast [DIC]), chloroplast division (autofluorescence [AF]), chloroplast division ring (FtsZ2-mNG), and cleavage furrow (EB1-mSc) structures.

**FIGURE 2:**
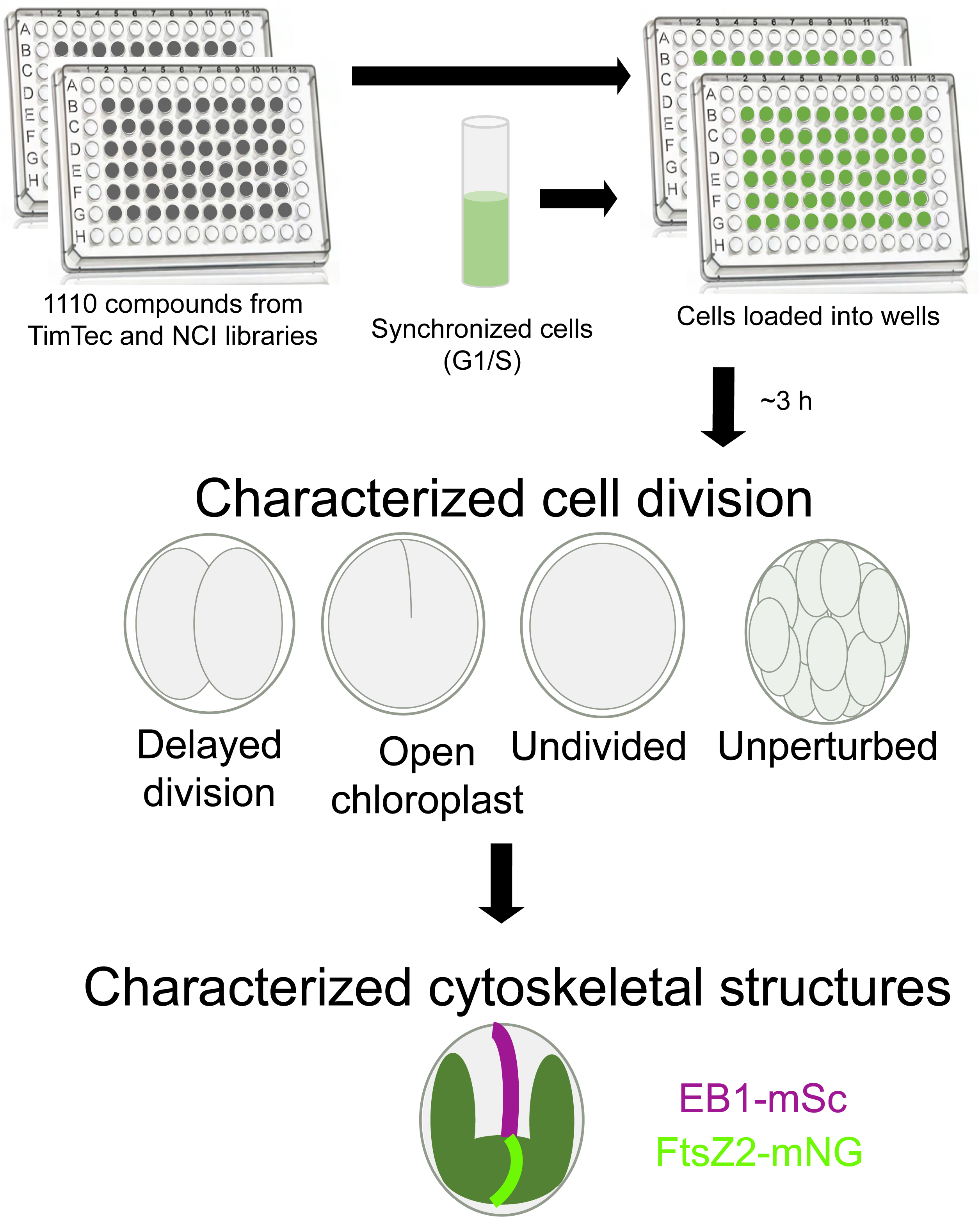
Overview of screen. Compounds were pre-loaded into 96-well plates; cells were synchronized in late G1 with alternating light-dark cycles and resuspended in TAP media; cell suspension was added to wells; after incubation, cell and chloroplast division was characterized; and where abnormal division was observed, FtsZ2-mNG and EB1-mSc structures were characterized.

Each compound produced some degree of heterogeneous effects, both within and among fields. To consolidate this variation into interpretable patterns, we first assigned a primary drug effect that reflected a field’s predominant appearance in seven categories: cell and chloroplast division, shape, size, and surface; microtubule structures; FtsZ2 structures; and the presence of vacuolation in the cytosol. Each field was assigned exactly one primary effect per category, and fields whose primary effect could not be clearly assigned for a given category were scored “uncategorized.” All primary effects for a single field were converted into effect frequencies for that hit, e.g., 40% of fields treated with epirubicin hydrochloride showed delayed division, 40% had open chloroplasts with notched cells, and 20% were undivided (Figure 3). Some fields were also assigned secondary effects, which were striking but not predominant (Figure S1). Finally, the 72 hits were classified into five groups based on K-modes clustering of the primary effects (see Materials and Methods) (Figure 3). Two compounds, curcumin and camptothecin, appeared in both NCI and TT libraries, for a total of 70 unique compounds (Figures 3 and S1). Of these 70 compounds, 67 have Chemical Abstracts Service (CAS) numbers with associated “type of compound” and/or “type of organism” term IDs. We identified terms that are significantly enriched in the 67 hits and in the individual K-mode clusters against the background of 693 compounds with CAS numbers that we screened (Figure 4A).

**FIGURE 3:**
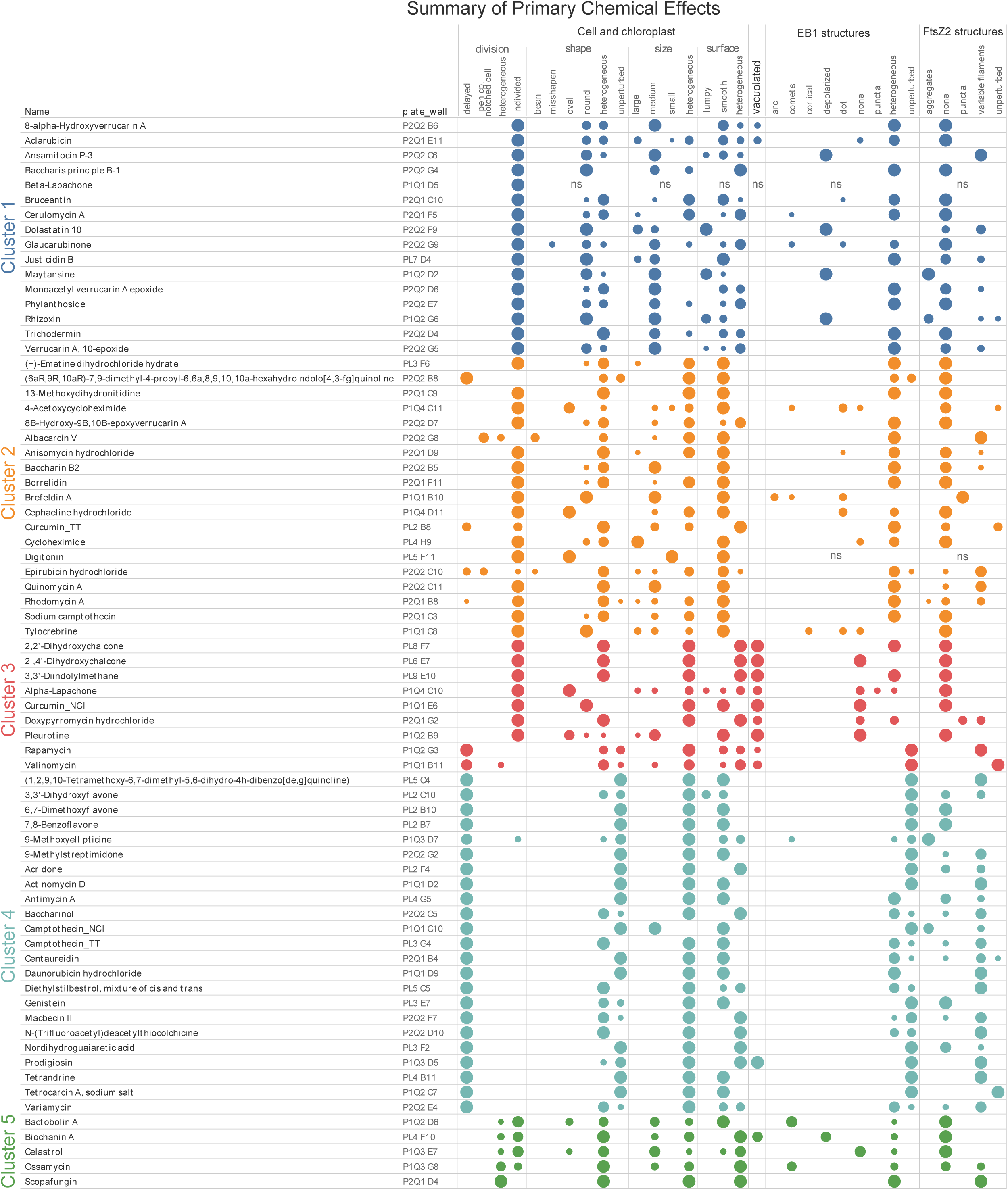
Summary of primary chemical effects for the 70 unique compounds (72 hits) that perturbed cell and chloroplast division. Treated cells were assessed for abnormalities in cell and chloroplast division, shape, size, and surface; vacuolation; EB1-mSc (microtubule) structures; and FtsZ2-mNG structures. Each colored circle (blue, orange, red, aqua, and green) represents a cluster, and the size of the circle corresponds to the frequency of that effect across all fields scored for that chemical. ns: not scored.

**FIGURE 4:**
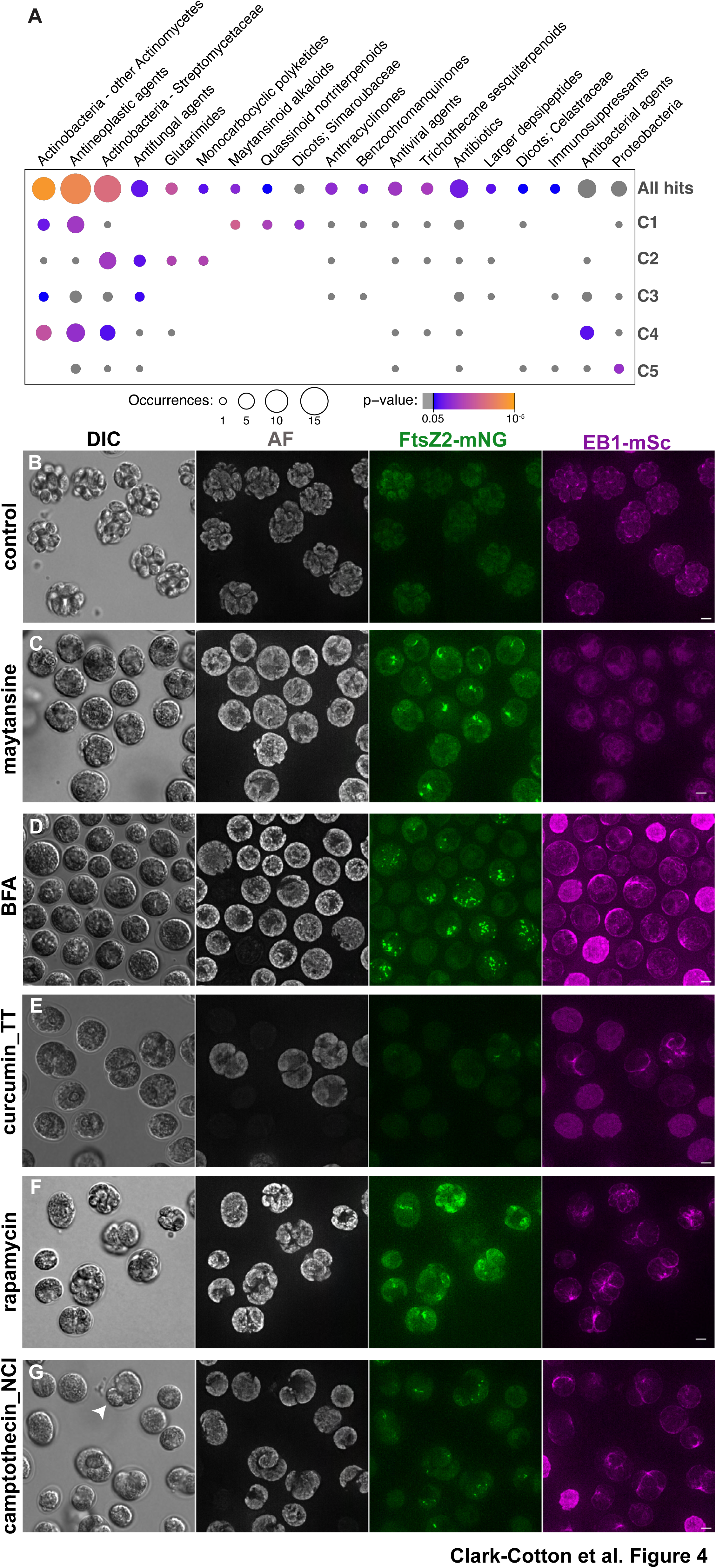
Examples of effects observed in the screen. (A) Enrichment analysis of chemical terms for the 72 hits (“All hits”) that disrupt cell or chloroplast division. Fisher’s exact tests were performed for existing chemical terms in the identified chemicals and five K-mode clusters against a background set (all examined chemicals in the screen). The colors indicate the p-values (grey, p ≥ 0.05), and the circle size represents the number of occurrences. No circle indicates zero occurrence of the term within the tested set. (B) In *nap1-1* cells treated with a non-perturbing chemical from the screening libraries, cells (DIC) and chloroplasts (AF) divided multiple times. At the time of imaging (3-5 hours after drug addition), no FtsZ2-mNG structures were visible in multiply divided cells, while indistinct EB1-mSc signals were observed. (B) When cells were treated with BFA, cells/chloroplasts did not divide. The FtsZ2-mNG signal appeared aggregated, and small polar arcs of EB1-mSc were sometimes visible. (C) Cells treated with camptothecin displayed delayed division or remained undivided, and asymmetrical division (arrow) was sometimes observed. Occasional indistinct FtsZ2-mNG signals were observed, and EB1 signals appeared to mark furrows in dividing cells. (D) Cells treated with curcumin also displayed delayed or no division phenotypes. FtsZ2-mNG signal was rare, but EB1-mSc appeared to mark cleavage furrows of rare dividing cells. Scale bars = 5 μm.

### Some division-perturbing compounds also alter cytoskeletal structures

Less than 7% of the 1110 compounds screened produced cell division effects. The remainder divided as expected, producing clustered daughter cells within the mother cell wall (Figure 4B). In these divided, unhatched cells, distinct FtsZ2 structures are absent, and EB1 appears polarized, as expected in post-division interphase cells.

Cluster 1 is highly enriched for antineoplastic agents (Figure 4A; aclarubicin, bruceantin, dolastatin 10, glaucarubinone, maytansine, and phyllanthoside). Cells treated with compounds in this cluster were all medium-sized and remained undivided, with the majority having no FtsZ signal, suggesting that the cells arrested growth before entering mitosis and forming an FtsZ ring (Figure 3). The exceptions are the four maytansinoid alkaloids (Figure 4A) that caused a complete loss of microtubules: ansamitocin P-3, dolastatin 10, maytansine, and rhizoxin, which were previously identified as microtubule polymerization inhibitors that bind the vinca binding site of tubulin (Ikeyama and Takeuchi, 1981; Bai *et al*., 1990). Cells treated with these compounds also showed persistence of FtsZ filaments or aggregates (Figs. 3 and 4C), suggesting that they had reached the cell-cycle stage at which FtsZ2 is expressed (normally in late G1) (Tulin and Cross, 2015; Zones *et al*., 2015), translocated into the chloroplast, and forms filaments (normally in pre-prophase, Figure 1A). These maytansinoid alkaloids may be useful alternatives to widely used microtubule inhibitors such as oryzalin and amiprophos-methyl (Collis and Weeks, 1978; James *et al*., 1988; Ehler and Dutcher, 1998).

Cluster 2 represents a wide variety of compounds that inhibit eukaryotic cellular processes (Figs. 3 and 4A), such as the glutarimides cycloheximide (CHX) and acetoxycycloheximide (ACH) (protein-synthesis inhibitors), the polyketides borrelidin (threonyl-tRNA synthetase inhibitor) and brefeldin A (BFA, an ER-Golgi transport inhibitor), and an antifungal agent, curcumin. Curiously, although CHX and ACH are analogs and both blocked cell division, CHX-treated cells grew to a large size, while ACH-treated cells were arrested with heterogeneous sizes (Figure 3). Because it has been reported that ACH has a higher toxicity than CHX for plants (and vice versa for fungi) (Nguyen *et al*., 2021), ACH may be a more potent inhibitor of protein synthesis in *C. reinhardtii*, especially at the low 25 µM concentration used in this study. BFA caused striking FtsZ2 structure abnormalities (Figure 4D): In addition to polarized localization of EB1-mSc either into a small polar cap or a larger arc, indicative of an early growth arrest due to secretion defects, FtsZ2-mNG appeared punctate or aggregated inside the chloroplasts. This result suggests that inhibition of the ER-to-Golgi transport (Helms and Rothman, 1992) has unexpected effects on the integrity of the FtsZ ring inside the chloroplast. Another compound in this cluster, curcumin (from the TT library), caused a near-complete loss of FtsZ2-mNG signal and failure in chloroplast division in some cells, while not blocking furrow ingression (Figure 4E; however, see below).

Cluster 3 is exemplified by vacuolation of the cells (Figure 3). Because of the static, non-time-lapse nature of the images screened, the timing of the formation of these vacuoles is unclear. Curcumin (from the NCI library) was identified for the second time with a more severe loss of FtsZ2 signal and no furrow formation. Rapamycin appeared to delay cell-cycle progression as reported previously (Jüppner *et al*., 2018), resulting in the formation of clusters of various numbers of cells at the end of the ∼3-h incubation period (Figure 4F).

All compounds in Cluster 4 caused delayed division, with the majority causing some forms of FtsZ2 filaments persisting in the chloroplast (Figure 3). This cluster is enriched for antibacterial agents (Figure 4A; actinomycin D, genistein, centaureidin, and nordihydroguaiaretic acid) which may target mechanisms of bacterial origin within the chloroplast. Among these, actinomycin D is a widely used transcription inhibitor that binds to DNA (Reich, 1963) and blocks gene expression from both the nuclear and plastid genomes in *Chlamydomonas* (Surzycki and Rochaix, 1971; Miller and McMahon, 1974; Guertin and Bellemare, 1979). It has been reported that actinomycin D blocks cell division as long as it is added to the cells before flagella resorption and DNA replication (Howell *et al*., 1975), which is consistent with our observation.

Camptothecin in this cluster (from both NCI and TT libraries) is an inhibitor of topoisomerase I involved in the regulation of DNA topology during replication, recombination, and transcription (Legarza and Yang, 2006). As previously reported (Voigt *et al*., 2017), cells treated with camptothecin showed division-phase-specific defects, suggesting that DNA replication in S phase is the major target (Figure 4G). Similarly to the effects of another drug that blocks the cell cycle in S phase (2-deoxyadenosine; Harper and John, 1986), camptothecin-treated cells did not enter mitosis yet aberrant furrows were formed, supporting the notion that initiation of mitosis is dependent upon completion of DNA replication, while initiation of cytokinesis is independent (Harper and John, 1986).

Cluster 5 represents a small number of compounds that showed large degrees of heterogeneity in most categories, including cell and chloroplast division. This likely reflects the general cytotoxicity of the compounds, such as a ribosome inhibitor, bactobolin A (Greenberg *et al*., 2020), and inhibitors of mitochondrial functions, ossamycin and scopafungin (Reusser, 1972; Salomon *et al*., 2000). Of note is that biochanin A is the fifth compound in this screen that showed depolymerized EB1 localization. Although biochanin A has not been previously reported to depolymerize microtubules, it is a derivative of genistein, which has been reported to inhibit microtubule polymerization (Mukherjee *et al*., 2010).

In summary, this screen identified putative inhibitors of cell and/or chloroplast division, many of which were previously unreported in *Chlamydomonas*. Because the screen was performed with fixed parameters (25 µM compound concentration, one time-point for imaging, one strain background), each of these compounds will require further validations.

### Curcumin inhibits FtsZ-ring formation and chloroplast division but not mitosis or cytokinesis

As a case study of validating compounds found in our screen, we focused on curcumin. In our screen, curcumin was included in both the NCI and TT libraries, and it produced slightly different results: While curcumin from NCI blocked both cell and chloroplast division with a complete loss of FtsZ2-mNG and EB1-mSc and vacuolation in some cells (Figure 3), curcumin from TimTec allowed some cells to form a partial cleavage furrow associated with EB1-mSc (Figs. 3 & 4E).

Curcumin is a chemical produced by turmeric and has been identified as a positive hit in various high-throughput screens (Ingólfsson *et al*., 2014). As a result, its use for the potential health and therapeutic benefits has caused some concerns and controversies (Liu *et al*., 2022). Molecularly, however, curcumin has been shown to directly bind to both FtsZ and tubulin. On one hand, curcumin binds to bacterial FtsZ proteins, accelerates GTP hydrolysis, inhibits protofilament formation and bundling in vitro (Rai *et al*., 2008; Kaur *et al*., 2010), and inhibits bacterial growth *in vivo*. On the other hand, curcumin binds to eukaryotic tubulin proteins, reduces their GTPase activity, and inhibits tubulin polymerization *in vitro* (Zhang and Kanakkanthara, 2020), and inhibits microtubule dynamics and cell proliferation *in vivo* (Gupta *et al*., 2006). The reported *K*_D_ values are higher for the curcumin-FtsZ interaction (∼7.3 µM) (Rai *et al*., 2008) than for the curcumin-tubulin (2.0-2.4 µM) (Gupta *et al*., 2006; Chakraborti *et al*., 2011). Our alignment of bacterial, archaeal, and plastidic FtsZ proteins showed that the curcumin-binding residues are largely conserved among these proteins (Figure 5A), suggesting that curcumin may bind to *Chlamydomonas* FtsZ1 and FtsZ2. Similarly, curcumin-binding sites on tubulins overlap with the vinca site that is conserved in *Chlamydomonas* (data not shown). Because *Chlamydomonas* simultaneously expresses tubulin and FtsZ proteins, it provides a unique opportunity to investigate the effects of curcumin on both systems within the same cell.

**FIGURE 5:**
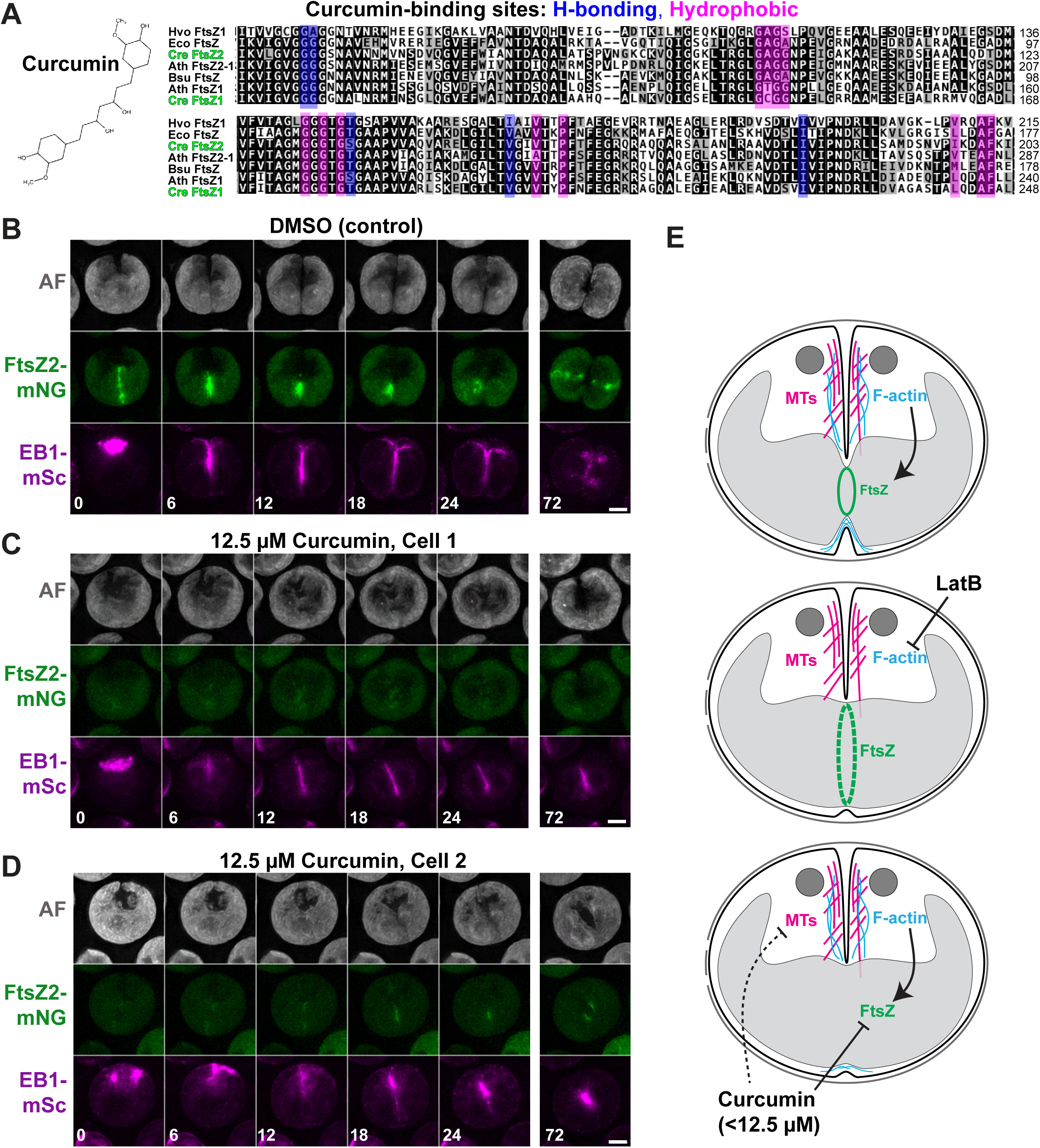
Effect of curcumin on FtsZ and chloroplast division. (A) The chemical structure of curcumin (left) and the conserved curcumin-binding residues in FtsZ homologs (right). Positions of the residues involved in hydrogen bonding (blue) and hydrophobic interactions (magenta) with curcumin in *in silico* docking of *Escherichia coli* and *Bacillus subtilis* FtsZ proteins are highlighted. Ath, *Arabidopsis thaliana*; Bsu, *B. subtilis*; Cre, *C. reinhardtii*; Eco, *E. coli*; Hvo, *Haloferax volcanii*. (B-D) Time-lapse images of SCC022 (FtsZ2-mNG EB1-mSc *nap1-1*) treated with DMSO or curcumin. (B) A control cell treated with 0.25% DMSO. An intact FtsZ2-mNG ring was formed before metaphase and constricted as the furrow ingressed. Both cell and chloroplast divided within 12-18 min. (CD) In two examples of cells treated with 12.5 µM curcumin (in 0.25% DMSO), FtsZ2-mNG was either (C) largely undetected or (D) appeared as weak signals at the division site with delay. In each case, a cleavage furrow was formed but did not complete division until the end of the imaging session (72 min). Scale bars = 5 μm. (E) A summary of the effects of LatB and curcumin on cell and chloroplast division. See main text for details.

We purchased curcumin from a commercial source and re-examined its effect on *Chlamydomonas.* At 25 µM used in the screen, localization of FtsZ2-mNG and EB1-mSc was completely abolished (Figure S2A), consistent with the results with curcumin from the NCI library. At a lower concentration of 12.5 µM (Figure S2A), FtsZ2-mNG failed to form a ring at the division site and instead formed puncta throughout the chloroplast; while many cells also lost EB1-mSc localization, some maintained seemingly normal EB1-mSc structures at the division site in dividing cells (Figure S2A, 12.5 µM, arrow). At 6.25 µM, many cells showed no or weak FtsZ2-mNG ring signal but with robust furrow-associated EB1-mSc (Figure S2A, 6.25 µM, arrows). Overall, curcumin appears to have a more substantial effect on FtsZ2 structures than on microtubules at lower concentrations. At all three concentrations, curcumin affected both FtsZ1-mNG and FtsZ2-mSc indistinguishably (Figure S2B), suggesting that this compound disrupts the entire FtsZ ring and not just FtsZ2.

The temporary disruption of the FtsZ ring by curcumin provides an opportunity to test the importance of coordinating cell and chloroplast division. Specifically, when chloroplast division is blocked, does the cell activate some checkpoint to delay the onset of cytokinesis, or does it still initiate cleavage-furrow ingression? If a furrow is formed, is it able to “cleave” the undivided chloroplast in the absence of a functional FtsZ ring? To test these, we performed time-lapse imaging of cells treated with 12.5 µM curcumin. Even in the presence of curcumin, cells were able to enter mitosis and form a spindle indistinguishable from control cells (Figure 5, B-D). Unlike control cells (Figure 5B; Movie S5), however, curcumin-treated cells did not have any FtsZ2 ring in metaphase (Figure 5, C and D, 0 min; Movies S6 and S7), although some cells had weak, transient FtsZ2-mNG signals in the middle of the chloroplast at later time points (see Figure 5D). Upon exit from mitosis, EB1-mSc in the curcumin-treated cells transitioned to the cell-division plane in a timely manner, suggesting that the onset of cytokinesis was not delayed (Figure 5, C and D; Movies S6 and S7). However, the chloroplasts did not divide before the end of time-lapse imaging, and EB1-mSc remained associated with the partially ingressed furrow (Figure 5, C and D; Movies S6 and S7). These results suggest that FtsZ-ring formation is not a prerequisite for normal mitosis and initiation of cytokinesis, but it is required for timely chloroplast division.

### Novel compounds to study cell and chloroplast division in *C. reinhardtii*

In this study, we examined over 1100 natural compounds isolated from various organisms for their effects on cell and chloroplast division in a green alga, *Chlamydomonas,* by a visual screen. Some of the hits are compounds previously known to inhibit cell growth and division in *Chlamydomonas* and/or other organisms (e.g., CHX, rapamycin, camptothecin), validating the approach. Some other hits provide new compounds targeting known pathways involved in cell division in *Chlamydomonas*, such as maytasinoids (microtubules) and ACH (protein synthesis).

Some hits caused somewhat perplexing observations. For example, BFA is an established inhibitor of ER-to-Golgi transport, yet it completely inhibited the formation of the FtsZ2-mNG ring in the chloroplast. In green algae and land plants, the chloroplast has two envelope membranes that are thought to be independent of the endomembrane system, and nuclear-encoded chloroplast-stromal proteins are translated in the cytosol and cross the envelopes through the translocons of the outer and inner chloroplast envelopes (TOC/TIC) (Nakai, 2018). Intriguingly, however, some glycosylated chloroplast-resident proteins in plants have been shown to go through what is termed the “ER-to-Golgi-to-plastid trafficking pathway,” and accumulation of such proteins is sensitive to BFA (Villarejo *et al*., 2005; Nanjo *et al*., 2006). Although FtsZ2 and other known chloroplast-division proteins have predicted signal peptides for direct TOC/TIC targeting and are not glycosylated, and thus are not likely to go through the ER-to-Golgi-to-plastid trafficking pathway, this pathway may transport some unknown protein(s) required for assembly of the FtsZ ring. If this pathway is indeed conserved*, Chlamydomonas* with BFA would provide a powerful research tool to understand its underpinnings.

In this study, we performed a further characterization of curcumin, a compound that has previously been shown to bind directly to tubulin and FtsZ. Our results suggest that, despite the higher affinity for tubulin *in vitro*, curcumin affects FtsZ ring formation at a lower dose *in vivo*. The reason for this apparent discrepancy is unknown, although it may be due to the difference in the molecular consequences of curcumin binding: curcumin accelerates GTP hydrolysis of FtsZ, which should promote protofilament disassembly, while it blocks GTP hydrolysis of tubulin, which should reduce the propensity for catastrophe once a microtubule is formed. Notwithstanding the molecular mechanism of inhibition, our results show that, at the low concentration range, curcumin inhibits FtsZ ring formation and chloroplast division while allowing for the cell to proceed through mitosis and cleavage-furrow ingression similar to the effects of loss of F-actin caused by LatB treatment (Figure 5E). These results suggest that F-actin may control the formation and/or function of the FtsZ ring through an unknown mechanism (Figure 5E). They also seem to suggest that the chloroplast-division checkpoint proposed in the red alga *Cyanidioschyzon merolae* may not be conserved in *Chlamydomonas.* In *C. merolae*, blockage of chloroplast division by overexpressing of FtsZ2-1 or a dominant-negative allele of dynamin-related protein 5B (Drp5B) caused a cell-cycle arrest in prophase with reduced cyclin B levels (Sumiya *et al*., 2016). Similar cell-cycle arrest upon chloroplast division failure was observed in the glaucophyte *Cyanophora paradoxa* (Sumiya *et al*., 2016). Because curcumin-treated *C. reinhardtii* cells with no FtsZ ring successfully entered mitosis and formed a cleavage furrow, it appears that the cell does not monitor whether the chloroplast is ready to divide. Alternatively, the FtsZ ring or protofilaments may be an essential component of the chloroplast-division checkpoint pathway, without which the checkpoint is no longer activated. A method to block chloroplast division with an intact FtsZ ring is required to dissect these possibilities. Regardless, it is noteworthy that land plants do not seem to have such a checkpoint, at least in the majority of their cell types that are multi-plastidic, given that their cells can tolerate inefficient chloroplast division caused by various defects including loss of FtsZ and DRP5B (Osteryoung and Pyke, 2014; Chen *et al*., 2018). It should also be noted that the apparent discrepancy may be due to technical reasons. For example, some of the defects observed in this study may involve subtle side effects of the curcumin on microtubules or other pathways, given the reported polypharmacological nature of this drug in general (Ingólfsson *et al*., 2014; Liu *et al*., 2022) and its dose-dependent effects on FtsZ and EB1 localization in *Chlamydomonas* in this study. Further validation using an *ftsz1Δ ftsz2Δ* mutant is required to access the side effects of curcumin, although such a mutant may be inviable if chloroplast division is completely deficient and blocks cell division. Conversely, curcumin and some of the compounds identified in this study may be effective in other organisms to perturb the cytoskeleton, providing additional avenues for interrogating the mechanisms of cytokinesis, chloroplast division, and their spatiotemporal coordination.

## MATERIALS AND METHODS

### Plasmids, strains, and growth conditions

Plasmid pMO699 (*P_EB1_:EB1-mScarlet:T_EB1_*:*Paro^R^*) was created by replacing *mNeonGreen* (an XhoI-XhoI fragment) in pEB1-mNeonGreen (Harris *et al*., 2016) with a PCR fragment encoding *mScarlet-I* from mScarlet-I-mTurquoise2 (a gift from Dorus Gadella; Addgene plasmid # 98839) by Gibson assembly (Gibson, 2009). Plasmid pMO773 (*P_FTSZ1_:FTSZ1-mNeonGreen-3FLAG:T_FTSZ1_*:*Hyg^R^*) was created by first inserting two PCR products, *P_FTSZ1_:FTSZ1-*EcoRV and EcoRV-*T_FTSZ1_*, amplified using genomic DNA as template into NotI-EcoRV-digested pRAM103 pRAM103 (Perlaza *et al*., 2019) by Gibson assembly. The resulting plasmid, pMO704 (*P_FTSZ1_:FTSZ1-HpaI:T_FTSZ1_*:*Hyg^R^*), was then digested with HpaI and assembled with a PCR product containing *mNeonGreen-3FLAG* from pMO665 (Onishi *et al*., 2020). Plasmids pMO774 (*P_FTSZ2_:FTSZ2-mNeonGreen-3FLAG:T_FTSZ2_*:*Hyg^R^*) and pSC018 (*P_FTSZ2_:FTSZ2-mScarlet_v2-PA:T_FTSZ2_*:*Hyg^R^*) were made with the same approach by first generating pMO705 (*P_FTSZ2_:FTSZ2-EcoRV:T_FTSZ2_*:*Hyg^R^*), then inserting a PCR product containing *mNeonGreen-3FLAG* from pMO665 or *mScarlet-I_v2-PA* from pRT113 (a gift from Ryutaro Tokutsu). These plasmids were linearized using appropriate restriction enzymes and transformed into *C. reinhardtii* by square-pulse electroporation (Onishi and Pringle, 2016). Transformants were selected for resistance to paromomycin (RPI, P11000; 10 µg/ml) or hygromycin B (VWR, 97064-454; 10 µg/ml) and subsequently for the expression of fluorescently tagged proteins.

*C. reinhardtii* wild-type strains CC-124 (mt-) and iso10 (mt+, congenic to CC-124; provided by S. Dutcher, Washington University in St. Louis, St. Louis) were the parental strains. Strain SCC022 (*nap1-1 EB1-mScarlet FtsZ2-mNeonGreen* mt-) used for screening was constructed as follows: Wild-type CC-124 (mt-) was transformed with *P_FTSZ2_:FTSZ2-mNeonGreen-3FLAG:T_FTSZ2_*:*Hyg^R^* (pMO774) and crossed with *nap1-1 mt+* to generate a *nap1-1 FTSZ2-mNeonGreen-3FLAG mt-* strain. This strain was then crossed with iso10 (*mt+*) that was separately transformed with *P_EB1_:EB1-mScarlet:T_EB1_*:*Paro^R^* (pMO699) to generate SCC022. Strain SCC059 (*nap1-1 EB1-mScarlet FtsZ2-mNeonGreen* mt-) was made by similar transformations using pMO773 and pSC018 and genetic crosses. The *nap1-1* mutant was previously isolated and backcrossed multiple times in the CC-124 background (Onishi *et al*., 2016).

All plasmids (with associated sequence files) and *C. reinhardtii* strains have been deposited to the Chlamydomonas Resource Center (https://www.chlamycollection.org).

Cells were grown in Tris-acetate-phosphate (TAP) liquid medium or TAP-agar plates (Gorman and Levine, 1965) at 25°C. For screening experiments, cells were synchronized with alternating 12-hour light:12 hours-dark (12L:12D) cycles (250 μmol m^2^ s^-1^) on TAP-agar plates for two days.

### Chemical-genetic screen

The TimTec Natural Product Library (720 compounds; timtec.net) and the National Cancer Institute Natural Products Set V (390 compounds; https://dtp.cancer.gov/organization/dscb/obtaining/available_plates.htm#nps_set) in DMSO were obtained in 96-well plates (10 mM, 0.25 ml per well, Greiner Bio-One SensoPlate^TM^ 96-well glass bottom microplates, #655892) from the Duke Functional Genomics Shared Resource.

Synchronized cells of strain SCC022 (*nap1-1 EB1-mScarlet FtsZ2-mNeonGreen* mt-) were collected in hour 9 of the light cycle and suspended in liquid TAP to a density of 1.0-2.0 × 10^^6^ cells/ml. At hour 10, 50 µl of the cell suspension was added to each well, followed by 50 µl of 1.5% low-melting-point agarose in TAP (55°C; IBI Scientific #IB70051) to immobilize the cells, to a final screening concentration of 25 µM of each compound. Plates were covered with lids and incubated at 25°C for at least 3 hours (2 hours of light and 1 hour of darkness) before imaging to optimize the opportunity to see altered division structures; imaging typically took ∼2h to complete (i.e., 3 hours into the dark phase).

From wells where cell division abnormalities were observed, DIC and fluorescence microscopy images were collected to reflect representative chemical effects on cell division (DIC), chloroplast division (autofluorescence), FtsZ2-mNeonGreen, and microtubules (EB1-mScarlet). Fluorescence microscopy was performed on a Leica Thunder inverted microscope equipped with an HC PL APO 63X/1.40 N.A. oil-immersion objective lens and an OkoLab incubator chamber that was maintained at 25-26°C. Signals were captured using following combinations of LED excitation and emission filters: 510 nm and 535/15 nm for FtsZ2-GFP (7%, 350 ms); 550 nm and 595/40 nm for EB1-mSc (5%, 100 ms); 640 nm and 705/72 nm for chlorophyll autofluorescence (AF; 1%, 10 ms), with 0.21 μm Z-spacing covering 8 μm. All images were processed through Thunder Large Volume Computational Clearing and Deconvolution (Leica). Fluorescence images were converted to maximum projections using ImageJ (https://imagej.net/software/fiji/), and the medial plane of DIC images was selected to produce single-plane images from each channel.

Two to eight images per channel were generally collected from each well, but due to transient microscope errors, there were three exceptions: plate P1Q1 well D5, from which only a single DIC image was collected; plate PL5 well F11, from which both DIC and fluorescence images were collected, but poor focus made the fluorescence images uninterpretable; and plate P2Q1 well D4, in which division was clearly abnormal but neither DIC nor fluorescence images were acquired. Images from all perturbing compounds, plus examples of unperturbed division used as controls, are available online (https://doi.org/10.6084/m9.figshare.28020959.v1).

### Screening scoring and analysis

Each screening image was visually inspected and assigned one code for each of seven primary categories, which reflected the predominant chemical effects observed in that field: cell/chloroplast division (delayed, open chloroplast/notched cell, heterogeneous, or undivided); cell/chloroplast shape (bean, misshapen, oval, round, heterogeneous, or unperturbed); cell/chloroplast size (large, medium, small, or heterogeneous); cell/chloroplast surface (lumpy, smooth, or heterogeneous); vacuolated (yes or no); EB1 structures (aggregates, arc, comets, cortical, depolarized, dot, none, puncta, heterogeneous, or unperturbed); and FtsZ2 structures (aggregates, none, puncta, rings, variable filaments, or unperturbed). Different images from a single well were scored independently, and the frequency of each effect’s occurrence was calculated for the well. When necessary, secondary effects were assigned to capture noteworthy observations that did not dominate the population, but only primary effects contributed to the frequency calculations. Two additional codes were required to score secondary effects on cell/chloroplast division: asymmetrical and open chloroplast. The 72 chemical compounds were clustered based on the scores in the seven primary categories by K-modes clustering (Huang, 1997) using the klaR package.

For the enrichment analysis of chemical terms, the chemical terms associated with the CAS registry numbers of each compound were extracted from the Dictionary of Natural Products of CHEMnetBASE (dnp.chemnetbase.com). Of the 1110 compounds screened, 692 had unique CAS registry numbers, and a “background” file of associated chemical terms was made. Input files of the same format were generated for six sets of chemicals: all identified hits from the screen (a total of 67 chemicals) and clusters C1 through C5 of the K-modes clustering. Next, a custom Python script was written to calculate the occurrences for each chemical term in the background and input files, and subsequently perform a Fisher’s exact test on a 2×2 contingency table for enrichment analysis of the term in the tested set relative to the background. The resulting datasets were filtered by removing duplicated chemical terms and terms with only one occurrence. The p-values and occurrences of the remaining 19 chemical terms across all identified hits and the 5 clusters were plotted as circular heatmaps using ggplot2 in RStudio.

### Time-lapse microscopy

For time-lapse imaging of cells treated with latrunculin B (Adipogen, AG-CN2-0031, Lot A01191/M), cells synchronized on TAP-agar were transferred to a TAP-agar with 3.0 μM LatB at hour 11 of the 12L:12D cycle. At hour 12.5, the cells on the TAP-agar + LatB plate were collected and placed onto an agarose block (1.5%, low-melting agarose in TAP-liquid) containing 3.0 μM LatB or 0.03% DMSO (control). The agarose blocks were placed down into wells of a chambered glass coverslip (Ibidi, 81817), and the additional space in between the agarose block and the edge of the well was sealed off by adding 2% low-melting agarose containing either LatB or DMSO. The time-lapse experiment started at hour 13, and images were captured at 5-min intervals for a duration of 1.5 hours.

For time-lapse imaging of curcumin (TCI, C2302), synchronized cells on TAP-agar plates were directly placed onto 1.5% agarose blocks containing different concentrations of curcumin at hour 11 of the 12L:12D cycle, and the time-lapse experiment started at hour 11.5 with 6-min intervals for a duration of 2.5 hours.

## ABBREVIATIONS

BFA: Brefeldin A
ACH: Acetoxycloheximide
CHX: Cycloheximide
LatB: Latrunculin B
NCI: National Cancer Institute
TT: TimTec

## ACKNOWLEDGMENTS

The authors thank Tavia Webley and Michael Cherry (American Chemical Society) and Sarah Shoemaker (North Carolina School of Science and Mathematics) for organizing summer high school student research opportunities. We thank Ryutaro Tokutsu for the gift of a Scarlet_v2 plasmid and So Young Kim of the Duke Functional Genomics Core Facility for assistance with procuring libraries. We also thank Eric Monson of the Duke Center for Data and Visualization Sciences for assistance with data visualization in Tableau.

This work was supported by NSF PRFB #2209112 to M.R.C-C, NSF #1818383 to J.R. Pringle and M.O., and NSF CAREER #2337141 to M.O. S-A.C. was a recipient of the Hung Taiwan-Duke University Fellowship. Support for A.G. and A.P. was provided by Project SEED of the American Chemical Society. Support for A.M. and V.M. was provided by the Mentorship Program at the North Carolina School of Science and Mathematics.

---

**FIGURE S1:**
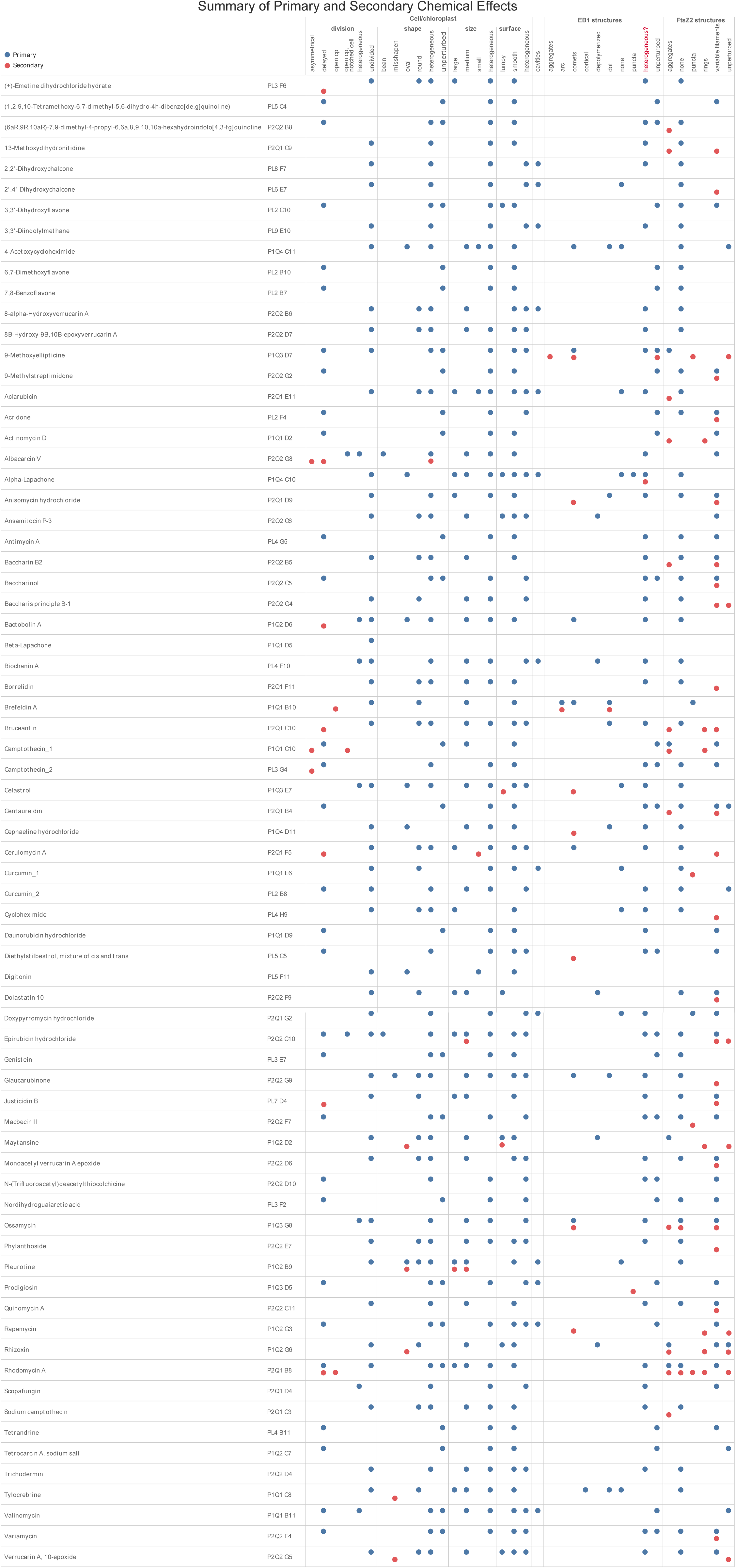
Summary of primary and secondary effects of the 70 unique compounds identified as perturbing cell and chloroplast division. All circles are the same size, regardless of the frequency of the effect. Blue: primary effects; red: secondary effects; ns: not scored.

**FIGURE S2:**
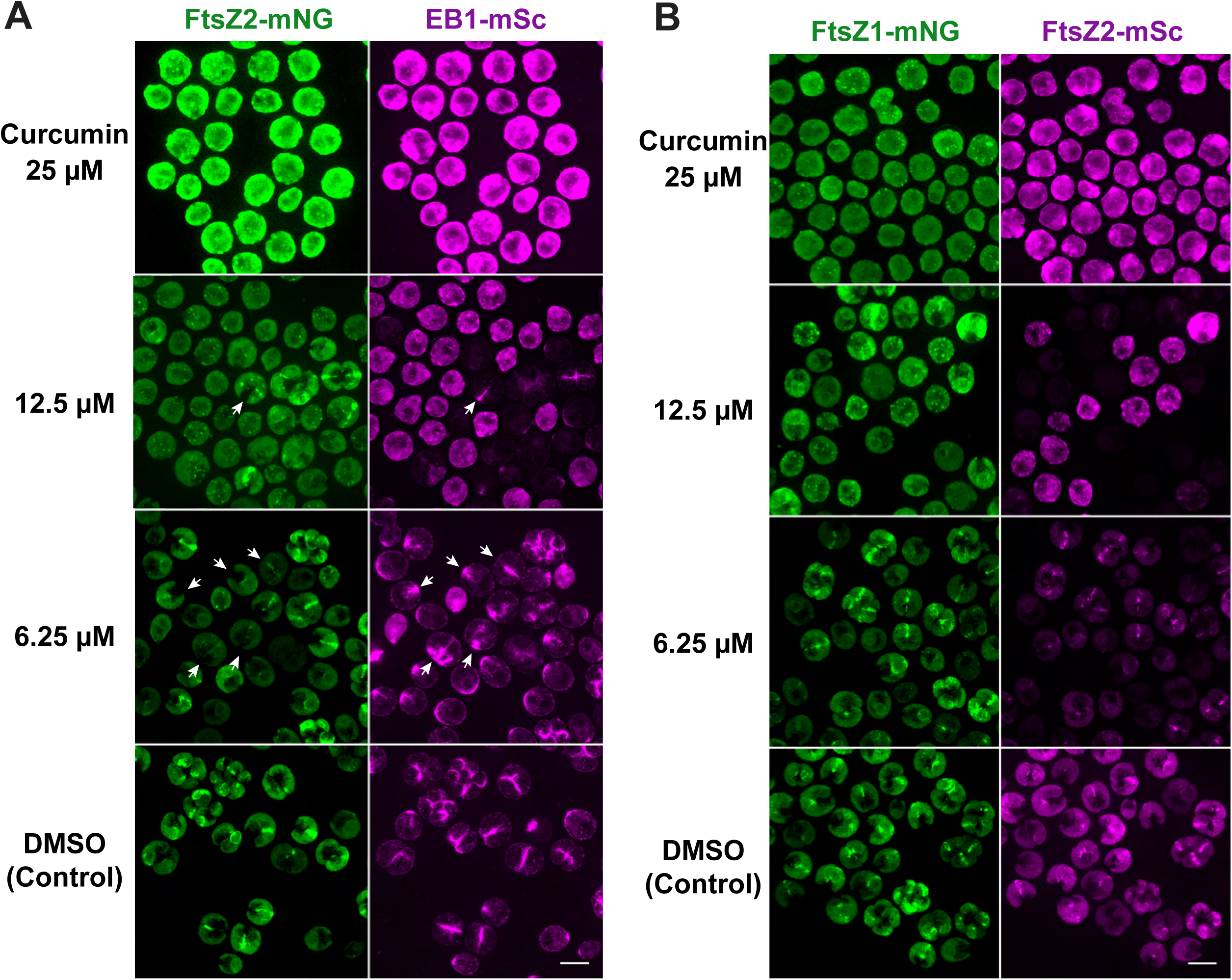
Time-course imaging of SCC022 (FtsZ2-mNG EB1-mSc *nap1-1*) and SCC059 (FtsZ1-mNG FtsZ2-mSc *nap1-1*) at different concentrations of curcumin. The cells were synchronized on TAP agar. At hour 11 of the 12L:12D cycle, the cells were moved to TAP agar containing 25, 12.5, and 6.25 μM curcumin, incubated at 21°C, and imaged at hour 13 (first hour into the dark period) by mounting the treated cells on TAP + 1.5% low-melting agarose with curcumin or DMSO (control, 0.25%). Arrowheads, cells with ingressing cleavage furrow associated EB1-mSc but no stable FtsZ2-mNG ring.

## Notes

### Competing Interest Statement

The authors have declared no competing interest.

